# A comparative study of endoderm differentiation in humans and chimpanzees

**DOI:** 10.1101/135442

**Authors:** Lauren E. Blake, Samantha M. Thomas, John D. Blischak, Chiaowen Joyce Hsiao, Claudia Chavarria, Marsha Myrthil, Yoav Gilad, Bryan J. Pavlovic

**Affiliations:** University of Chicago, Department of Human Genetics, Chicago, IL; University of Chicago, Department of Medicine, Chicago, IL

**Keywords:** Comparative genomics, functional genomics, gene expression

## Abstract

There is substantial interest in the evolutionary forces that shaped the regulatory framework that is established in early human development. Progress in this area has been slow because it is difficult to obtain relevant biological samples. Inducible pluripotent stem cells (iPSCs) provide the ability to establish in vitro models of early human and non-human primate developmental stages. Using matched iPSC panels from humans and chimpanzees, we comparatively characterized gene regulatory changes through a four-day timecourse differentiation of iPSCs (day 0) into primary streak (day 1), endoderm progenitors (day 2), and definitive endoderm (day 3). As might be expected, we found that differentiation stage is the major driver of variation in gene expression levels, followed by species. We identified thousands of differentially expressed genes between humans and chimpanzees in each differentiation stage. Yet, when we considered gene-specific dynamic regulatory trajectories throughout the timecourse, we found that 75‥ of genes, including nearly all known endoderm developmental markers, have similar trajectories in the two species. Interestingly, we observed a marked reduction of both intra-and inter-species variation in gene expression levels in primitive streak samples compared to the iPSCs, with a recovery of regulatory variation in endoderm progenitors. The reduction of variation in gene expression levels at a specific developmental stage, paired with overall high degree of conservation of temporal gene regulation, is consistent with the dynamics of developmental canalization. Overall, we conclude that endoderm development in iPSC-based models are highly conserved and canalized between humans and our closest evolutionary relative.

## Introduction

Differences in gene regulation between humans and other primates likely underlie the molecular basis for many human-specific traits (*1*). For example, it has been hypothesized that human-specific gene expression patterns in the brain might underlie functional, developmental, and perhaps cognitive differences between humans and other apes (*2, 3*). A recent comparative study that explored the temporal dynamics of gene regulation found potential differences in the timing of gene expression in the developing brain across primates (*4*). The authors argued that such differences might be related to inter-species differences in the timing of developmental processes. More generally, comparative studies in primates, while challenging, have already resulted in a few important insights into the evolution of gene expression levels and the traits they are associated with (*5*). Yet, we are also finding that gene expression patterns alone (without additional context or perturbation) provide little insight into adaptive phenotypes, molecular mechanisms, or even the specific biological processes involved in the observed changes in gene expression levels.

The challenge is that comparative studies in humans and non-human apes are extremely restricted because we only have access to a few types of cell lines and to a limited collection of frozen tissues (*5*). A few studies have chosen to sidestep this limitation by using model organisms in an attempt to recapitulate inter-primate differences in gene regulation. In this approach, the phenotypic impact of differences in gene regulation between humans and non-human primates are studied with high spatial and temporal resolution in a model species, such as the mouse (*6-9*). These studies are useful and often informative, but when model organisms are used to recapitulate gene regulatory differences between primates, the inference about function requires one to make a critically important assumption, which is typically not tested. Namely, one must assume that the effects of gene regulatory changes on complex phenotypes are identical in model organisms and in humans. This is a common assumption, not exclusively made in the context of comparative studies. Indeed, every study of human disease that makes use of model organisms relies on this assumption to a certain extent. The difference is that model system studies of human disease ultimately have to seek evidence that inference based on model systems is relevant to humans. In contrast, studies of human evolution using model systems rarely, if at all, are required to meet this standard. Indeed, oftentimes it is unclear how to design an experiment in model organisms that can directly address a phenotype difference between humans and non-human primates, for instance, when the phenotypes under consideration are related to cognitive abilities. In such cases, the assumption that the effects of gene regulatory changes on complex phenotypes are identical in model organisms and in humans is strained (*8*).

The caveats associated with using a model organism to study the phenotypic effects of regulatory differences between primates notwithstanding, until recently, it is not clear that there was an alternative approach. Indeed, while current comparative studies using primate material (tissue samples or cell lines) have provided valuable insight into the genetic architecture of gene regulation, we did not have a flexible and faithful framework with which to to dynamically study inter-species variation in gene regulation (*5*). In particular, frozen post-mortem tissues are not optimal templates for many functional genomic assays; as a result, we lack datasets that survey multiple dimensions of gene regulatory mechanisms and phenotypes from the same individuals (*5, 10*). Moreover, because it is rare to collect a large number of tissue samples from the same donor (particularly in non-human primates), we have never had the opportunity to study cross-species, population-level patterns of gene regulation in multiple tissues or cell types derived from the same genotype (same donor). We also have not been able to comparatively study population-level dynamics of gene regulation in primates, for example, during perturbation. In order to gain true insight into regulatory processes that underlie variation in complex phenotypes, we must have access to faithful model systems for a wide range of tissues and cell types. In other words, to utilize comparative functional approaches to comparatively study the genetic architecture of complex phenotypes in humans and other apes, a new approach is needed.

Recent technological developments in the generation and differentiation of induced pluripotent stem cells (iPSCs) now provide a renewable, staged and experimentally pliable source of terminally differentiated cells. Utilizing timecourse differentiation protocols, we can examine the context dependent nature of gene regulation, as well as the temporal roles of gene expression as different cell types and developmental states are established (*11*). This approach seems promising, and indeed, a handful of recent studies have been successful in utilizing iPSCs from humans and chimpanzees to characterize the uniquely human aspects of craniofacial development (*12*) and cortex development (*13, 14*).

Primate iPSC panels are a particularly attractive system for comparative studies of early development. With recent advances in iPSC technology and sequencing, we can now begin to examine whether specific phases of development undergo similar levels of constraint in primates. To this end, we chose to differentiate iPSCs from human and chimpanzee into the endoderm germ layer; from which essential structures in the respiratory and digestive tracts are ultimately derived, including the liver, the pancreas, and the gall bladder, the lung, the thyroid, the bladder, the prostate, most of the pharynx and the lining of the auditory canals and the larynx (*15*). Using this system, we found evidence to support developmental canalization of gene regulation in both species, 24 hours after differentiation from an iPSC state.

## Results

### Study design and data collection in the iPSC-based system

To perform a comparative study of differentiated cells, we used a panel of 6 human and

4 chimpanzee iPSC lines previously derived and characterized by our lab (*16, 17*). We differentiated the iPSCs into definitive endoderm, a process that was completed over 3 days (*11*), and included replicates of cell lines that were independently differentiated (see Methods and Figure 1a). We assessed the purity of the differentiated cells on each day using flow cytometry with a panel of six canonical markers, corresponding to the cell types we expected in the different stages of differentiation (Figure S1; Supplementary Data S1).

**Figure 1.**
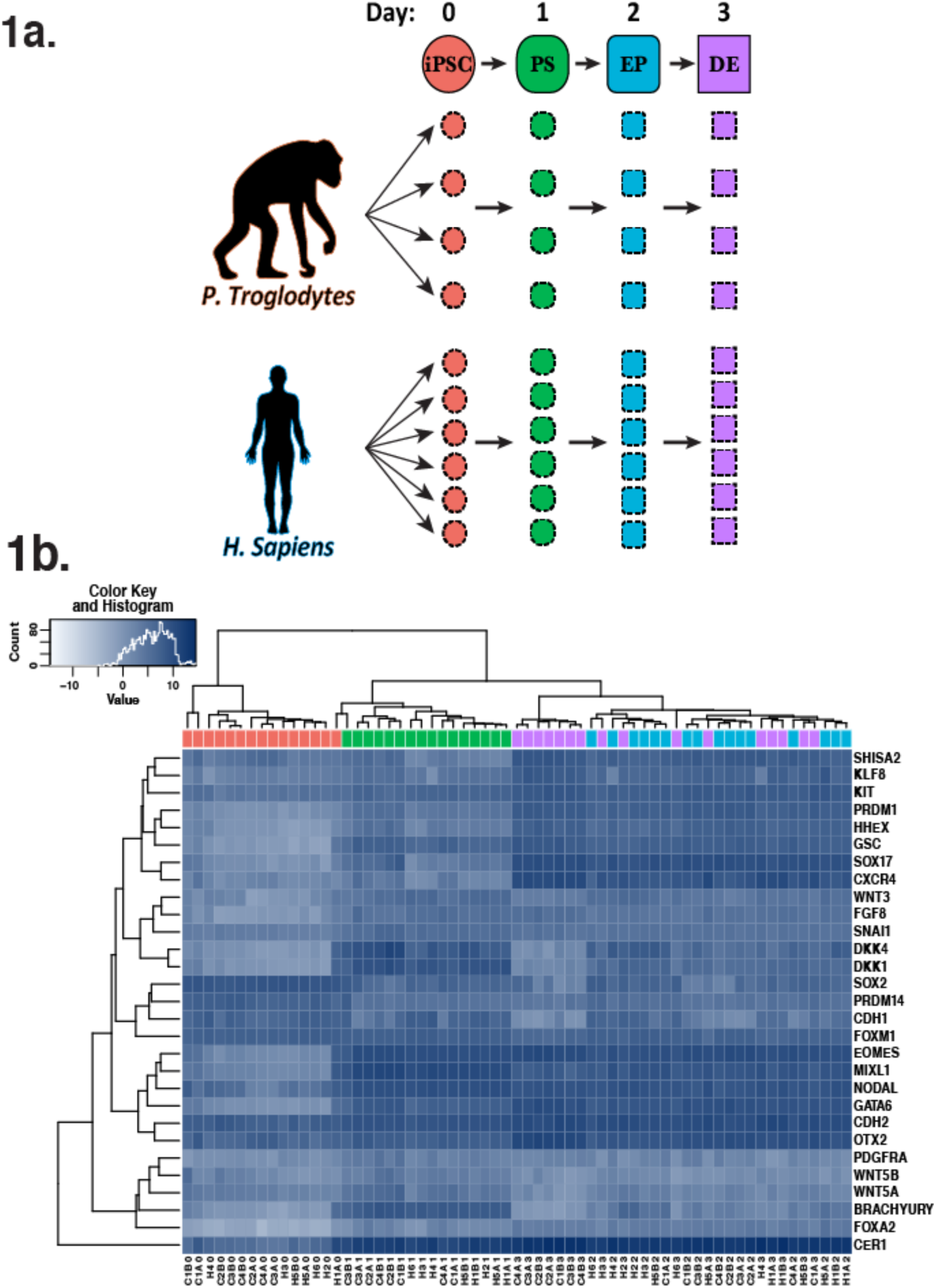
**a.** Study design. Four chimpanzees and six humans were studied at four time-points during endoderm development. (Two technical replicates from each of the chimpanzees and two technical replicates for two of the six humans for a total of 16 samples per time point.) iPSC: induced pluripotent stem cell, PS: primitive streak, EP: endoderm progenitor, DE: definitive endoderm. **b.** Heat map of normalized log_2_(CPM) as a measure of expression levels of transcription factors that are known to be highly expressed in one or more stages in the differentiation to endoderm (*11*). Generally, samples from the same day, regardless of species, cluster together.

We also harvested RNA from iPSCs (day 0) prior to differentiation and subsequently every 24 hours to capture intermediate cell populations corresponding to primitive streak (day 1), endoderm progenitors (day 2), and definitive endoderm (day 3). Overall, we collected a total of 32 human samples and 32 chimpanzee samples (Figure 1a). We confirmed that RNA from all samples was of high quality (Figure S2; Supplementary Data S2) and subjected the RNA to sequencing to estimate gene expression levels. Detailed descriptions of all individual donors, iPSC lines, sample processing and quality, and sequencing yield, can be found in the Methods section and Supplementary Data S3.

To estimate gene expression levels, we mapped reads to the corresponding genome (hg19 for humans and panTro3 for chimpanzees) and discarded reads that did not map uniquely (*18*). We then mapped the reads to a list of previously described metaexons across 30,030 Ensembl genes with one-to-one orthology between human and chimpanzee (*10, 19*). We eliminated genes that were lowly expressed in either species and normalized the read counts using the weighted trimmed mean of M-values (TMM) algorithm and a cyclic loess normalization by species and day, within individuals (Figure S3) (*20, 21*). We removed data from one clear outlier sample (H1B at Day 0; Figure S3) and repeated this process with the data from remaining samples to obtain TMM-and cyclic loess-normalized log_2_ counts per million (CPM) values for 10,304 orthologous genes (Figure 2a; Figure S4a; Supplementary Data S4). These normalized gene expression values were used in all downstream analyses. To mitigate the potential effect of gene length differences between the species on expression estimates using RNA sequencing, we also calculated reads per kilobase of orthologous exonic sequence per million mapped reads (RPKM) from the normalized read counts (Figure S4b).

**Figure 2.**
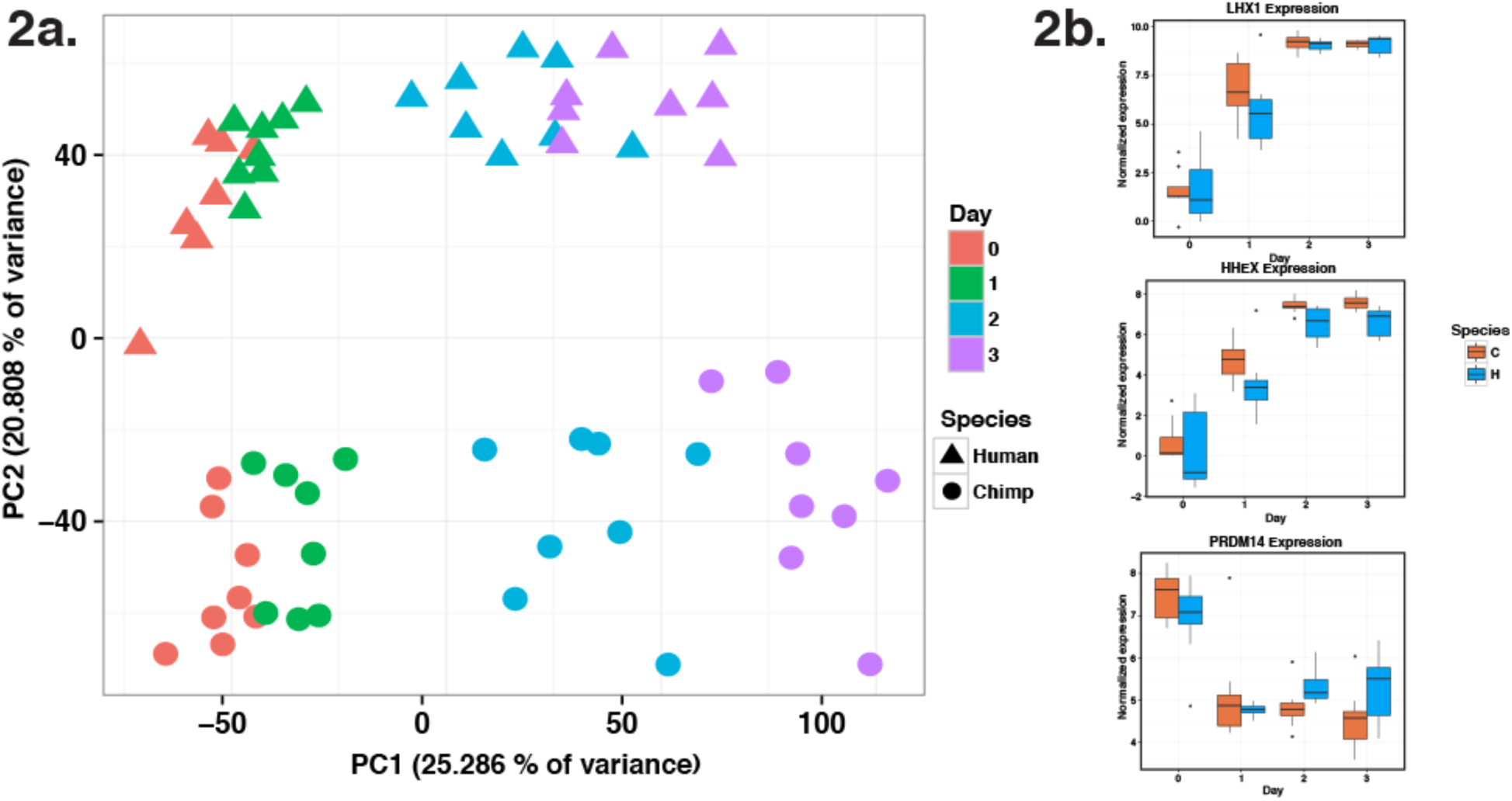
**a.** Normalized log_2_(CPM) expression measurements for all genes projected onto the axes of the first two principal components. Color indicates day. Shape represents species. PC1 is highly correlated with differentiation day (r = 0.92). PC2 is highly correlated with species (r = 0.93). **b.** Three box plots of normalized expression values for genes with known roles in endoderm development.

### iPSCs-based system effectively models primate endoderm differentiation

A global survey of the gene expression data (using principal component analysis) indicated that the primary sources of gene expression variation are differentiation day (Figure 2a; Supplementary Data S5; regression of PC1 of normalized gene expression levels by differentiation day, *P* < 10^-15^), followed by species (regression of PC2 of normalized gene expression levels by species, *P* < 10^-15^). This observation was also supported by clustering analysis based on the correlation matrix of pairwise comparisons of the gene expression levels (Figure S5).

Given the potential impact of study design properties on gene expression data and subsequent conclusions (*22*), we confirmed that none of our recorded variables related to sample processing were confounded with our main variables of interest, namely day and species (Supplementary Data S3, S5). We note that when all sequencing pools (mastermixes) were considered together, there was a relationship between adaptor sequence and day (χ^2^ test, Benjamini-Hochberg adjusted *P* = 0.01); however, this relationship is substantially weaker when ‘adaptor sequence’ and ‘day’ were tested in each of the 4 sequencing pools separately (B.H. adj. *P* > 0.9 in each test). Our most highly dependent variables with day or species were related to properties inherent to the iPSC model, including harvest density and day (B.H. adj. *P* = 0.01), harvest density and species (B.H. adj. *P* = 0.03), and harvest time and day (B.H. adj. *P* = 0.01) (Figure S6; Supplementary Data S3, S5). Overall, we were confident that our study design was

After characterizing global gene expression patterns, we focused on the expression of specific transcription factors with known roles in developmental pathways (Figure 1b) and other previously known lineage specific markers (*11, 23, 24*). Consistent with the results of our FACS analysis (Figure S1), we observed that the temporal trajectory of expression levels of known lineage specific markers and transcription factors further supported the assumed differentiation stages in each day (e.g. primitive steak-specific markers had increased expression on day 1, Figure 2b). The lineage specific markers and transcription factors were expressed at comparable levels in humans and chimpanzees at the relevant time points, consistent with previous literature (*11, 23*), and further supporting the validity of our *in vitro* system (Figures 2b, S9b).

### Comparative assessment of gene expression changes during differentiation

To identify gene expression differences between humans and chimpanzees throughout the timecourse, we used the framework of linear models (see Methods). We first assessed how many genes were differentially expression (DE) across species at each time point independently. Using this approach, we observed that the number of DE genes between humans and chimpanzee was similar across all time points (at FDR of 5‥ we classified 4475 – 5077 genes as DE at the different times points; Figure 3a; Supplementary Data S6). Nearly half of the genes that were classified as DE between the species in any single time point were found to be DE in all time points (2269 genes, 22‥). Nearly a third of genes whose expression was measured in our experiment were not classified as DE between the species at any time point (2862 genes, 28‥). Taken together, these observations suggested a strong relationship (violating the assumption of independent sampling) between gene regulatory patterns across samples from the different differentiation days, as expected.

**Figure 3.**
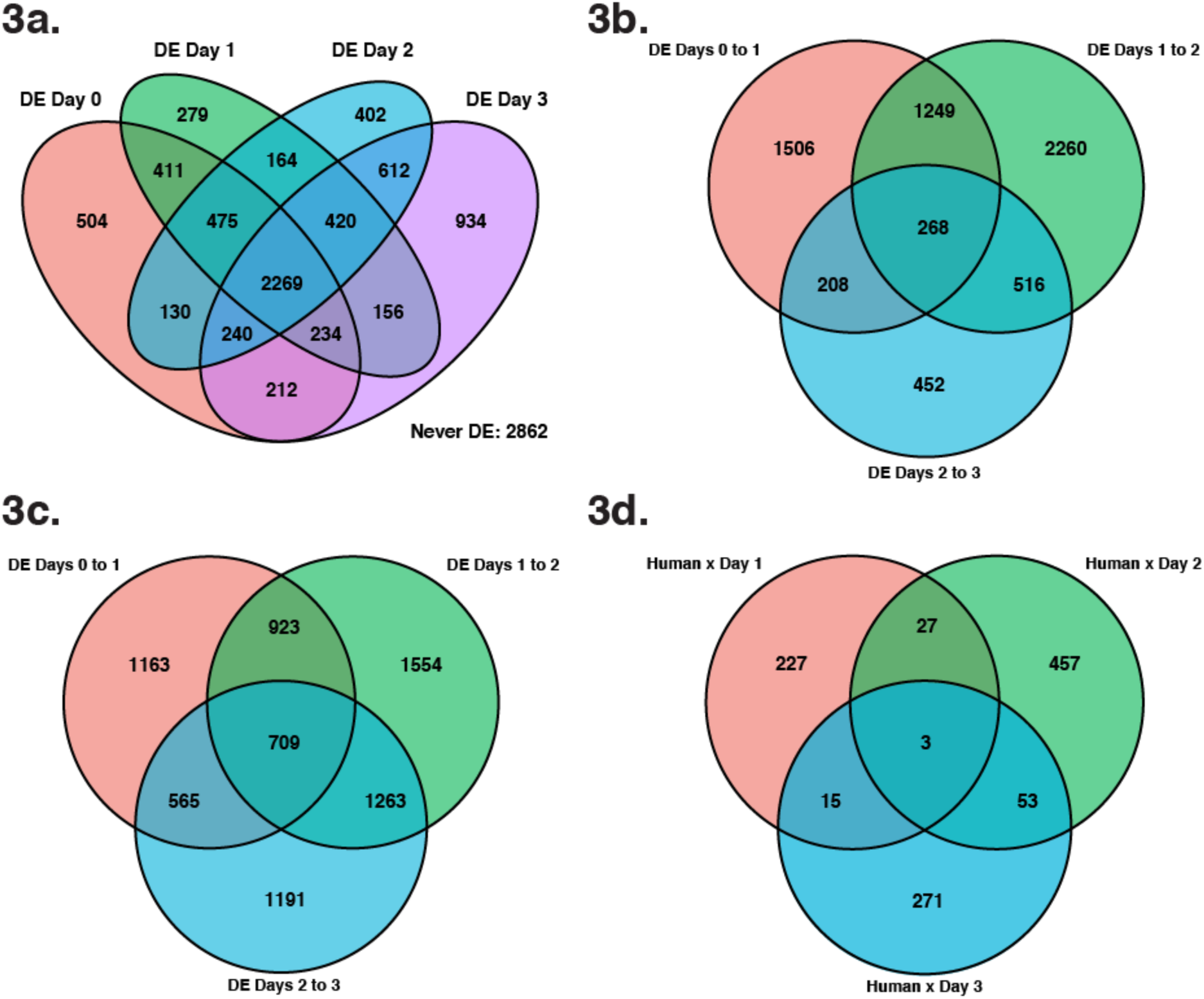
Number of differentially expressed (DE) genes in pairwise analyses. a. Venn diagram of all DE genes at each day (5‥ FDR). **b.** Venn diagram of all DE genes between consecutive time points in humans (5‥ FDR). **c.** Venn diagram of all DE genes between consecutive time points in humans (5‥ FDR). **d.** Venn diagram of genes with a significant species-time point interaction effect at each day (5‥ FDR).

We proceeded to consider temporal expression patterns within species. We considered expression changes across consecutive time points and found that we have more power to detect temporal gene expression differences in chimpanzee compared to humans (Figures 3b, 3c, 4; Supplementary Data S7), especially with respect to the transition between endoderm progenitors (day 2), and definitive endoderm (day 3). Despite this potential technical difference between the species (see Discussion), we found substantial evidence for conservation of temporal regulatory patterns in early differentiation states, with a possible increased divergence in definitive endoderm. Indeed, when we accounted for incomplete power (see Methods), we estimated that 77‥ of DE genes between iPSCs and primitive streak in humans are also DE between these states in chimpanzees; that 77‥ of DE genes between primitive streak and endoderm progenitors in humans are also DE between these states in chimpanzees; and that 80‥ of DE genes between endoderm progenitors and definitive endoderm in humans are also DE between these states in chimpanzees (Supplementary Data S8). As might be expected from these observations, we found that the relationship between day and gene expression was largely independent of species (Figure 3d; Supplementary Data S9).

### Joint Bayesian analysis reveals conservation of temporal gene expression profiles

In an attempt to overcome issues of incomplete power affecting these original naïve pairwise DE comparisons, and to account for dependency in data from different time points, we utilized a Bayesian clustering approach implemented by Cormotif (*25*). This joint modeling technique leverages expression information shared across time points to identify the most common temporal expression patterns (referred to as “correlation motifs”).

We identified diverse expression patterns that emerge as differentiation progresses in both species (Figure 5, motifs 2-8; Supplementary Data S10) as well as a set of 3789 genes whose expression is not significantly altered throughout the timecourse (8004 genes could be reliably classified into a motif, see Methods for inclusion criteria; Figure 5, motif 1; Supplementary Data S10). We found further evidence for conserved gene expression patterns early in the timecourse, as 75‥ of genes assigned to a motif were assigned to motifs with the same (Figure 5, motif 5) or similar temporal regulatory trajectories in both species (Figure 5, motifs 2, 3, 8). When we discounted the definitive endoderm samples, where we suspect that a technical confounder has increased variance between the species, we assigned 85‥ of genes to motifs with the same temporal trajectories across species (Figure 5, motifs 3, 5, 6-8; see Discussion). These observations are robust with respect to the number of correlation motifs (Figure S8b, Supplementary Information), the method used to combine data from technical replicates (Figure S8c), which days were included in the pairwise comparisons (Figure S8d), and the inclusion of all 10,304 genes in the analysis (Supplementary Information). Our Cormotif results are also consistent with the degree of conservation in gene expression trajectories that we obtained by using a correlation-based method to analyze relative changes in gene expression through the timecourse (Supplementary Information, Figure S9). Finally, the result that regulatory divergence in definitive endoderm is highest in our study is also constent with the results of the linear model based framework.

We found two correlation motifs with a potential marked difference between the species at a given stage (Figure 5, motif 4 with 187 genes and motif 7 with 686 genes). In both of these motifs, data from the earliest time points were conserved but gene regulation in the final stage (day 2 to 3) differed between the species. The genes in these motifs were enriched for Gene Ontology (GO) annotations related to animal organ development (e.g. *NRTN*, *PITX2*, *RDH10*), anatomical structure morphogenesis (*ARHGDIA, EHD2, SERPINE1*), regulation of developmental process (*FLRT3*, *LOXL2*, *SEMA7A*), and regulation of cell differentiation (*DIXDC1, ENC1, IRF1*; Bonferroni corrected *P* < 0.029 for all of these GO annotations in the PANTHER Overrepresentation Test; complete results in Supplementary Data S11) (*26-28*). In contrast, these four GO annotations were not enriched in other similarly sized motifs (Figure 5, motif 2) or group of motifs (Figure 5, motifs 3, 5).

### Reduced variation in gene expression levels at primitive streak

We next turned our attention to differences in the magnitude of variation in gene expression levels across time points. Previous studies reported that variation in gene expression levels between individuals was lower in iPSCs than in differentiated cells including lymphoblastoid cell lines (LCLs) and iPSC-derived cell types (*29*). We were thus interested in gene expression variation during iPSC differentiation.

We first compared within-species expression variation for all 10,304 orthologous genes across time points. Considering the distribution of expression variation across all genes, we found a marked reduction in inter-individual variation of gene expression levels as the human samples differentiated from iPSCs to primitive streak (*P* < 10^-15^, Figure 6; see Methods). We also detected this pattern when we considered the chimpanzee samples (*P* < 10^-15^, Figure 6), but the effect size in chimpanzee is much smaller. The differences in effect sizes notwithstanding, we did not identify similar reduction in variation in gene expression levels in any other transition during the timecourse in either species (*P* > 0.5 for testing the null of no change gene expression variance from day 1 to 2 and from day 2 to 3 in each species).

**Figure 6.**
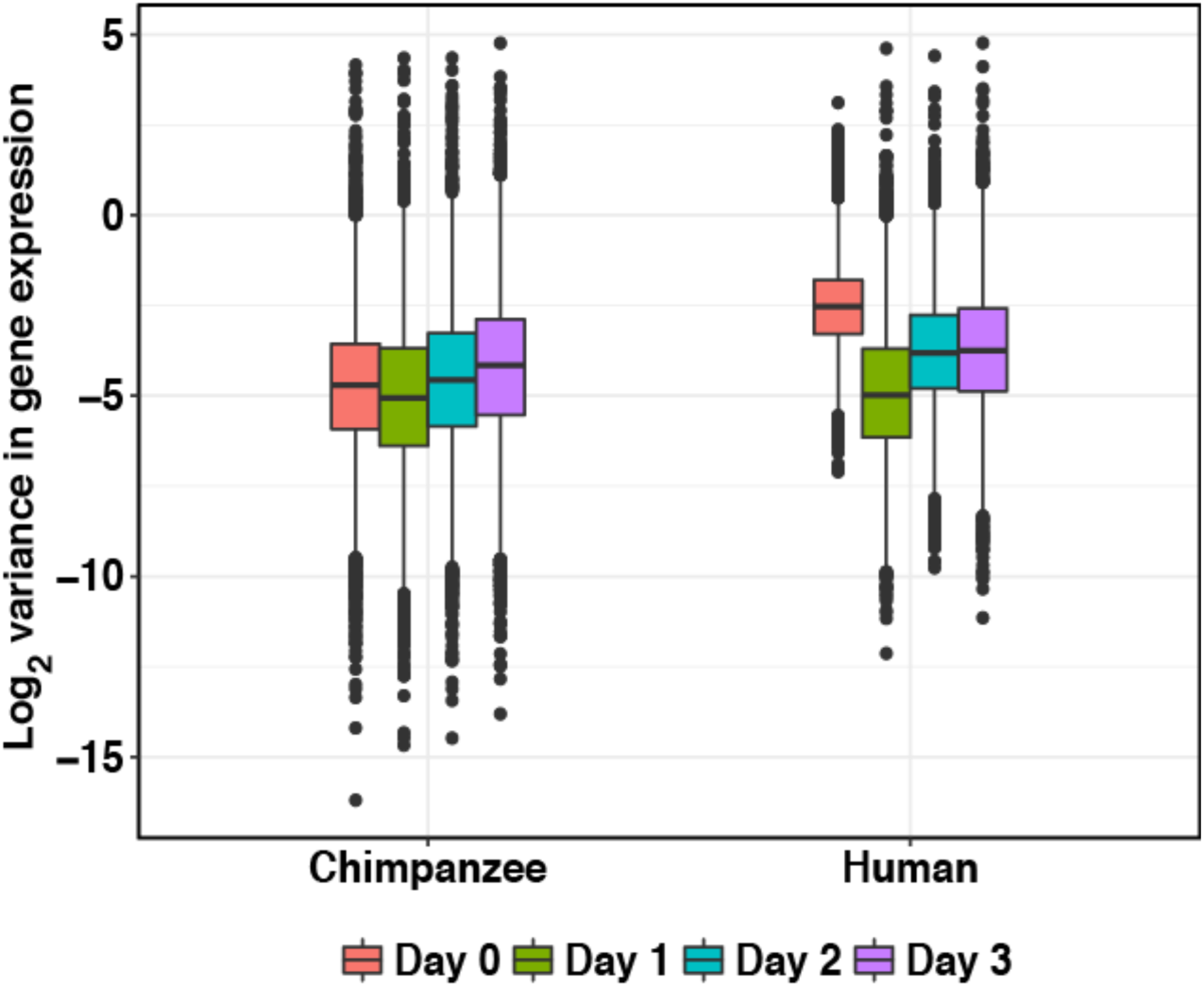
Global reduction of variation in gene expression from the iPSCs to primitive streak state. Box plot of the log_2_ variances of gene expression levels for each gene. Variation in gene expression levels are significantly reduced from iPSCs to primitive streak (*P* < 10^-15^ in both species) but not in subsequent time points (*P* > 0.5 in both species).

The overall human-chimpanzee divergence in gene expression levels was also slightly reduced as samples differentiated from iPSCs to primitive streak (Figure S10, Mann-Whitney *U* Test, *P* = 0.04), but not in any other transition during the timecourse. Furthermore, while we classified 504 genes as DE between humans and chimpanzees exclusively in iPSCs (of a total of 4,475 DE genes in iPSCs; FDR = 5‥, Figure 3a, Supplementary Data S12), we found only 279 genes that were DE exclusively in primitive streak samples (from a total of 4,408 DE genes for the primitive streak). The number of genes that are DE between the species exclusively in endoderm progenitors and definitive endoderm samples is higher (at FDR of 5‥, 402 and 934, respectively, Supplementary Data S12). The difference in the number of DE genes between these differentiated states could potentially be explained by a number of non-biological factors. Nevertheless, this observation is intriguing given that within species variation in gene expression levels – especially in humans -is lowest in primitive streak samples (namely, given a reduced variance, one would intuitively expect to have more power to detect inter-species DE genes between primitive streak samples).

The observation of a smaller number of genes that are DE exclusively in primitive streak samples compared with iPSCs is robust with respect to the FDR cutoff, differentiation batch, and purity of the samples (Figure S11; Supplementary Data S12). We also determined that the recorded technical factors are highly similar across biological conditions in days 0 and 1, and therefore are not likely to explain this observation (see Methods; Supplementary Data S5, S13). We thus proceeded to analyze the trajectory of variation in expression level on an individual gene basis. In this analysis, we were particularly interested whether the individual genes that undergo a change in variation of expression levels are shared to both species or not.

We used F tests to identify genes whose within-species variation in expression levels differs across time points (see Methods). Distributions of *P* values from all tests can be found in Figures 7a-b and Supplementary Data S14, which indicate that for a large number of genes, within-species variation in expression levels were reduced specifically and exclusively in primitive streak samples. Indeed, while we did not have much power to detect differences in variation in individual gene expression levels between states (due to such a small number of individuals in each species) we observed a clear excess of small *P* values from day 0 to 1, indicating a departure from the null expectation. Using Storey’s approach (*30*) to account for incomplete power, we estimated that within-species variation in expression levels was reduced as the samples differentiate from iPSCs to primitive streak in 83‥ and 27‥ of human and chimpanzee genes, respectively (Figure 7a-b, Day 0 to 1; see Methods). This result was robust with respect to the method used to calculate the proportion of true positives (*31*) (Figure S12).

**Figure 7.**
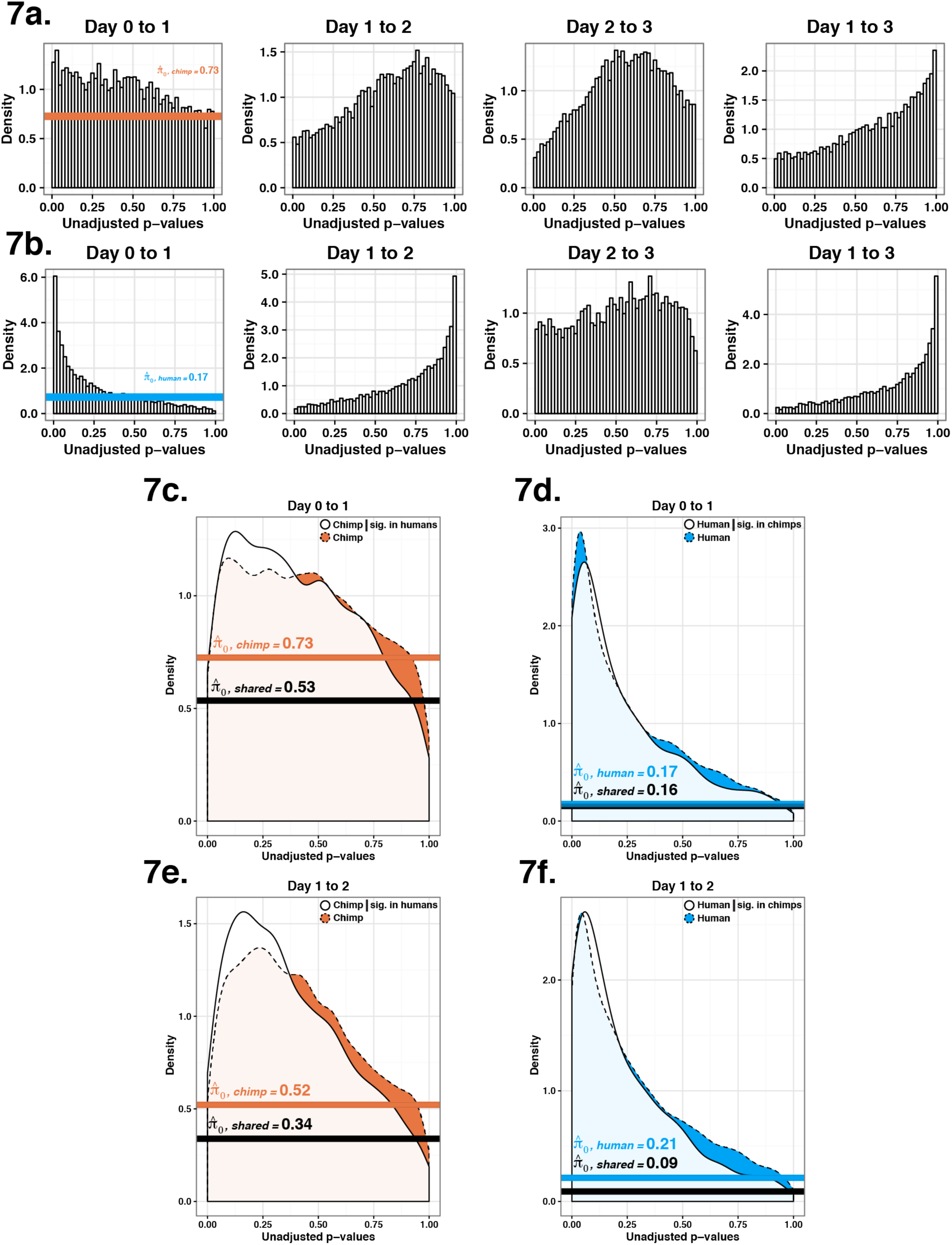
Genes with reduced variation in gene expression values at the primitive streak often demonstrate this property in both species. a. *P* value distributions from F tests of the null that variation in gene expression has not changed in chimpanzee samples. p0 is the proportion of null statistics in each distribution, so lower p0 values correspond to a larger estimate of true positives. **b.** *P* value distributions from F tests of the null that variation in gene expression has not changed in human samples. **c.** *P* value distribution from Figure 7a (orange) compared to the *P* value distribution calculated when one only considers genes for which reduced variation has already been observed (P < 0.05, white). **d.** Same as Figure 7c but the *P* value distribution was generated using human samples (blue) conditioned on significant genes in the chimpanzee samples (P < 0.05, white). **e.** *P* value distribution from an F test considering gene expression data from days 1 and 2 in the chimpanzee samples (orange) compared to the chimpanzee samples conditioned on the human samples (P < 0.05, white). **f.** *P* value distribution from an F test considering gene expression data from days 1 and 2 in the human samples (blue) and in the human samples conditioned on the chimpanzee samples (P < 0.05, white).

The proportion of genes with reduced within-species variation in expression levels in primitive streak samples is quite different between humans and chimpanzees (Figure 7a-b, Day 1 to 2). Yet, we had not observed this pattern in any other differentiation state in our data (Figure 7a-b, Days 1 to 2, 2 to 3, and 1 to 2). We thus asked about the overlap of genes with reduced variation in primitive streak samples across the two species. Specifically, we asked whether human genes with lower within-species variation in expression levels in primitive streak are more likely to show the same pattern in chimpanzee genes. For this analysis, we again used the Storey approach (*30*) to estimate the proportion of true positive tests in one species, conditional on the observation of reduced variation in the other species (using a relaxed cutoff of unadjusted *P* value of 5‥; see Methods). Using this approach, we found evidence that the pattern of reduced within-species variation in gene expression levels in primitive streak samples is generally conserved. We estimated that 47‥ of genes whose variation in expression level is reduced in human primitive streak samples showed a similar pattern in chimpanzees (under a permuted null we expect 27‥, *P* < 10^-4^, see Methods; Figures 7c, S13a; Supplementary Data S15). When we condition on observing a reduction of variation in chimpanzees, the overlap with humans was 84‥ (this high value was not unexpected because of the initial large proportion of human genes with a clear signature of reduced variation in primitive streak; under a permuted null we expect 83‥, *P =* 0.38; Figures 7d, S13b; Supplementary Data S15).

Using a similar approach, we also found a marked overlap of genes whose expression underwent a significant increase in variation throughout the transition from primitive streak to endoderm progenitors (Figures 7e, Π0 = 0.34 in chimpanzee genes conditioned on those with previously observed increased variation in the humans; Figure 7f, Π0 = 0.09 in human genes conditioned on those with previously observed increased variation in the chimpanzees). All our observations were robust to a wide range of statistical cutoffs used to classify genes whose within-species variation changes across the differentiation states (Figure S14).

## Discussion

In our opinion, the most significant finding of this study is the observation that regulatory variation is markedly reduced in both humans and chimpanzees, as the cell cultures differentiate from iPSCs to primitive streak. We believe that our finding that regulatory trajectories throughout endoderm differentiation are generally highly conserved in these two species was expected. Yet, our observation that a large number of genes are associated with reduced regulatory variation in a specific transition state, in both species, is a somewhat surprising property.

Before we discuss the potential implications of our observation, we first highlight study design considerations for our iPSC-based differentiation models and a few caveats. Since we designed the study to facilitate cross-species comparisons, we used human and chimpanzee iPSC lines that were generated, and then differentiated, using the same protocols. We made considerable efforts to balance the majority of sample processing properties related to our study design with respect to species and time point. For example, the two differentiation batches we used included multiple human and chimpanzee lines (which we also balanced with respect to gender). We included a number of technical replicates across the batches, but we were able to include 4 replicates for chimpanzees and only 2 for humans. The greater number of chimpanzee technical replicates may have contributed to increased precision and thus to power to detect regulatory differences during the timecourse in chimpanzees.

*In vitro* differentiation protocols are not identical to natural developmental signaling. Natural cell-driven developmental processes may be overridden by our administered media conditions. Indeed, even the most effective *in vitro* differentiation protocols are less effective than natural *in vivo* developmental signaling pathways. Thus, we used flow cytometry analysis with specific markers to estimate the purity of cell cultures at all times points in the second differentiation batch (Figure S1). Although we used an identical differentiation protocol across the entire experiment, we observed a wide distribution of cell purity across samples. In most time points the distribution of cell purity was not associated with species (this was reassuring), but in the last day, as samples differentiated to definitive endoderm, we observed a clear difference in cell purity between species (Figure S1d; Supplementary Data S1). As far as we can determine, the potentially technical inter-species difference in definitive endoderm purity should have no impact on our conclusions with respect to regulatory patterns in earlier differentiation time points.

More generally, we cannot exclude the possibility that small differences in cellular heterogeneity can explain our observations, at least in part. Cellular heterogeneity has been explored in developmental studies as a potential factor that drives cell fate determination. For example, a study of single-cell gene expression during the first four days of mouse development revealed a progressive increase in variability in expression level of lineage specifying transcription factors (TFs) from blastomeres in the 8 cell stage to cells in the inner cell mass. The higher variability in the timing of TF expression changes in the inner cell mass is thought to underlie fate decisions between primitive endoderm and epiblast commitment (*32*). Similarly, a single cell gene expression study in human pre-implantation embryos has proposed that heterogeneity in the downregulation of genes in human blastomeres contributes to lineage predispositions (*33*). A recent study of single cell gene expression during differentiation of embryonic stem cells down each of the three lineages primary germ layers (endoderm, mesoderm, and ectoderm), revealed that cells differentiating down the endoderm lineage exhibit the most asynchrony, indicating that there is likely substantial heterogeneity of degree of differentiation at any given time point (*34*).

Differences in cellular heterogeneity across time points may have driven the observation of reduced regulatory variation in primitive streak samples. In our opinion this is unlikely, because we expect the iPSC cultures to be the most homogenous in our experiment; nevertheless, we cannot provide data to exclude this explanation. In other words, without corresponding single cell gene expression data, it is difficult for us to distinguish between canalization of regulatory trajectories and reduced cellular heterogeneity as potential explanations for our observations. Single cell RNA sequencing data will also be able to shed light on our observation that gene regulation in definitive endoderm (day 3), unlike earlier days, does not indicate strong conservation between the species. At definitive endoderm samples, we observed the largest number of DE genes across species (Figure 3a; Supplementary Data 6), the smallest number of DE genes between two consecutive time points in humans (Figure 3b; Supplementary Data 7), and the lowest overlap in DE genes between time points (Figure 4). Unlike in the human samples, the chimpanzees had a relatively consistent number of DE genes between any two consecutive time points (Figure 3c; Supplementary Data 8). Some of these observations might be explained by the cell purity difference between species.

**Figure 4.**
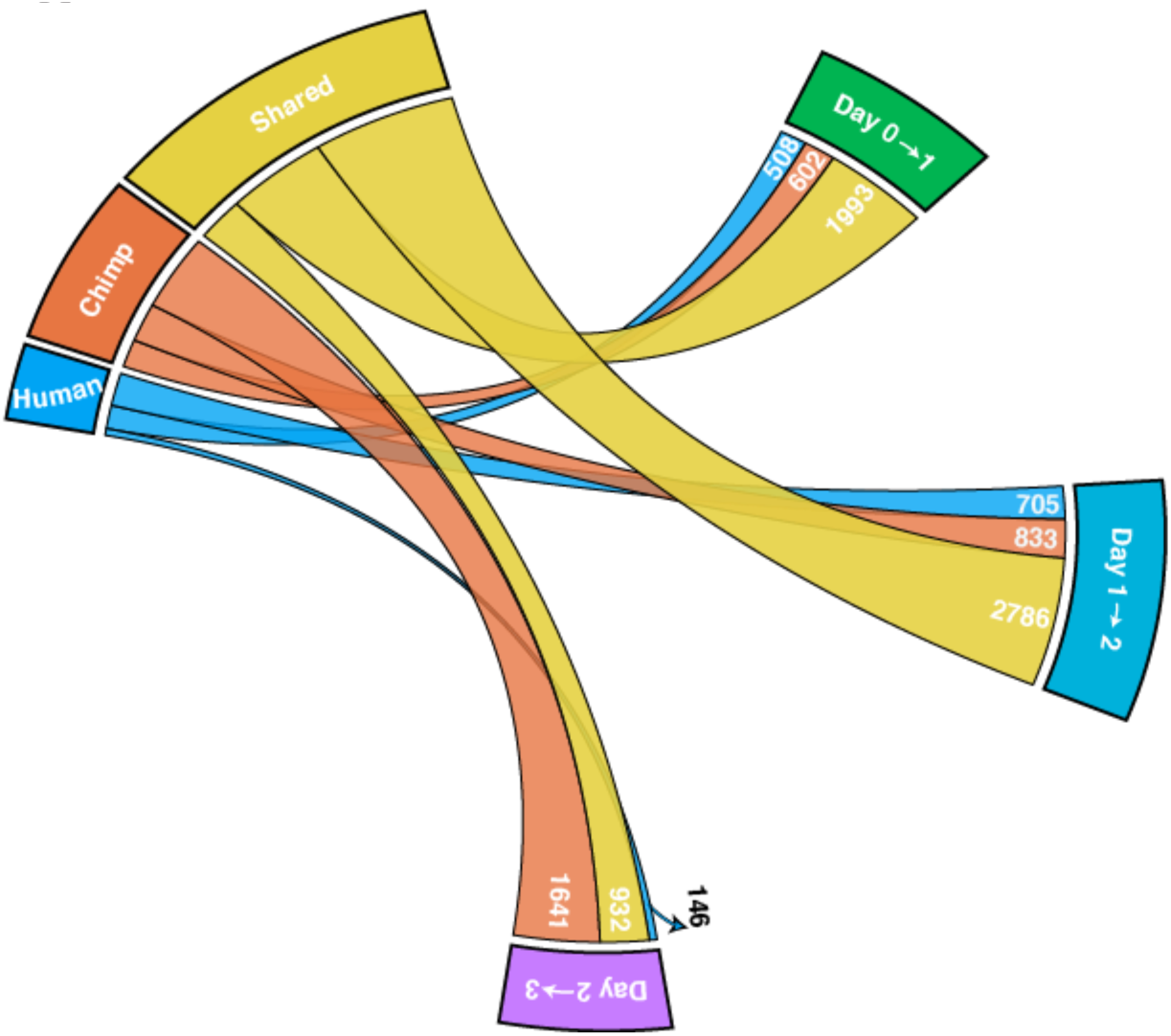
A circos diagram with the number of shared, human-specific, and chimpanzee-specific DE genes across time points. There is a high degree of sharing of DE genes (yellow ribbon), particularly from day 0 to 1 and day 1 to 2.

There is clearly a need for greater resolution, both temporal and spatial, in future studies of endoderm differentiation. Nevertheless, we argue that our observations indicate a strongly conserved temporal expression profile across species during early differentiation. Two lines of evidence support this conclusion. First, we observed a large number of genes with similar expression profiles across species. Indeed, we found that DE genes between differentiation states are shared between the two species far beyond what expected by chance alone (Supplementary Data S8). Our observations likely underestimated the proportion of shared regulatory patterns due to incomplete power, and consequently, our inability to reject the null should not be interpreted as strong support for the null. Moreover, when we jointly analyzed data from the entire timecourse, nearly all the gene expression trajectory motifs we identified, including 75‥ of all genes assigned to a motif (Figure 5), are shared across the two species. (The proportion of genes in the same gene expression trajectory motifs across species rises to 85‥ if we exclude data from definitive endoderm). Second, the observation of reduced variation of gene expression within and across species at the primitive streak stage supports a highly regulated and conserved process.

**Figure 5.**
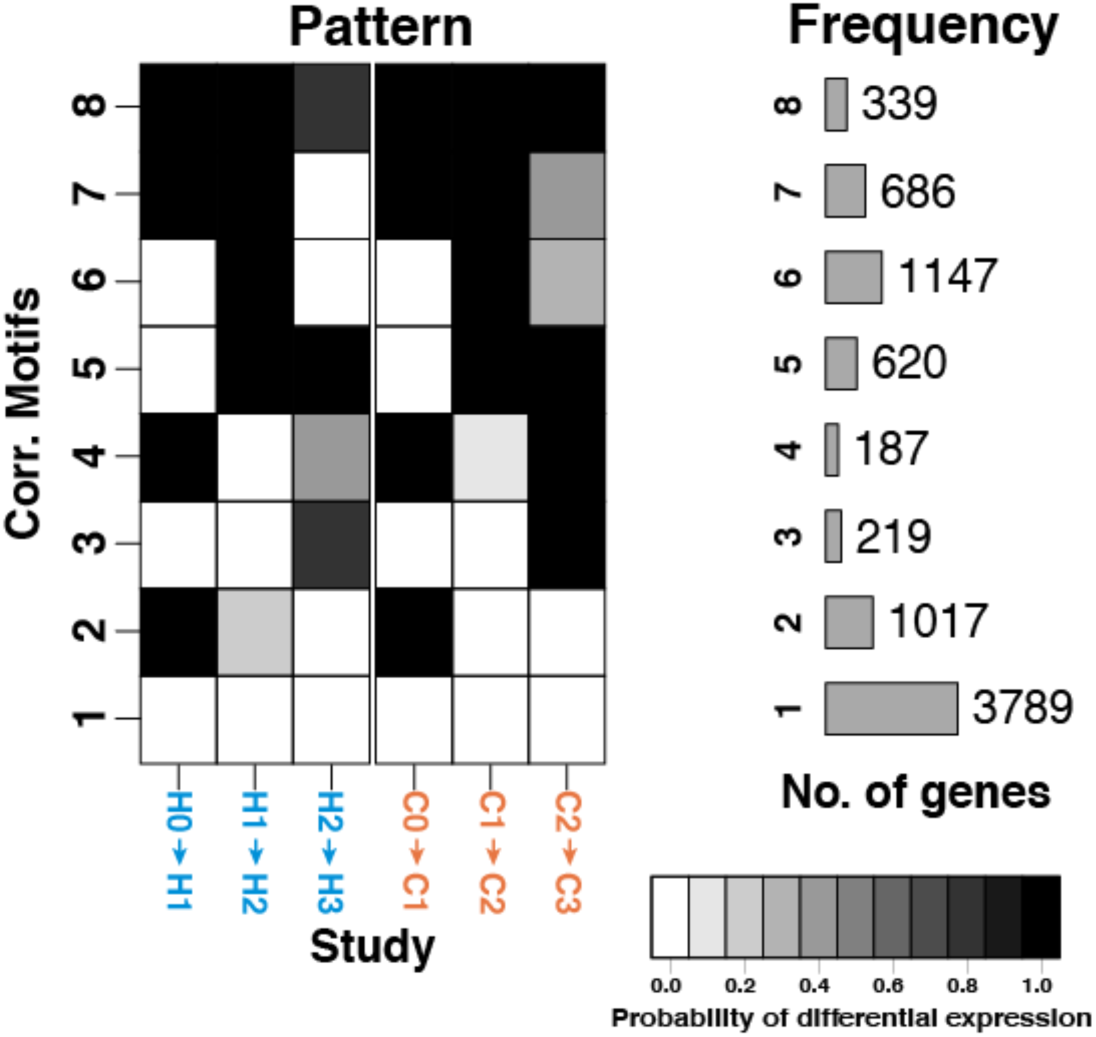
Correlation motifs with the probability of differential expression across days for each species and the estimated number of genes belonging to each correlation motif. The shading of each box represents the posterior probability that a gene is DE between two time points in a given species. Each row (“correlation motif”) represents the most prevalent expression patterns along the trajectory. 8004 out of the 10,304 total genes were assigned to one correlation motif in this model.

The observation of reduced regulatory variation is rather unusual in general, partly due to the unusual design of our study. Indeed, only few comparative studies have been designed to allow one to measure changes in variation over time. One such example, from a different context, can be found in a previous study in which monocytes from humans, chimpanzees, and rhesus macaques, which were stimulated with lipopolysaccharide (LPS) to mimic infection (*35*). When comparing gene expression in LPS-stimulated monocytes to that of non-stimulated cells, the authors found a reduction of inter-species variation in gene expression levels in a number of key transcription factors involved in the regulation of *TLR4*-dependent pathways. (It should be noted that this study did not focus on within-species variation changes.) In our current study, we found a more general change in regulatory variation across differentiation states, within and across species, without partitioning the genes into particular pathways or networks. Perhaps the number of regulatory pathways involved in an early developmental lineage commitment is be higher than those involved in a cellular response to stimulus by lineage committed cells, or perhaps – as has been suggested previously (*36*)-developmental pathways need to be more tightly regulated in general.

Indeed, reduced regulatory variation early in the endoderm differentiation process may be driven by the property of canalization during development. The theory of canalization posits that developmental processes end in a finite number of states despite minor environmental perturbations (*36-38*). Canalization is fundamentally linked to evolutionary states (*37*), and thus phenotypic robustness; therefore, even when reduced variation in gene expression levels is observed in cell culture, the explanation of canalization is intuitively appealing considering the discrete nature of cell types in an adult animal. Our results suggest that stages subsequent to primitive streak may follow a more relaxed transcriptional regulation with higher influence of individual genotypes. Our observations, therefore, may be consistent with activation of deeply conserved regulatory programs at the initial stages of gastrulation followed by processes less affected by evolutionary constraint and therefore potentially more amenable to adaptation. In other words, our results supports the expectation that gastrulation is a highly canalized and conserved process in humans and chimpanzees.

More generally, we believe that despite limitations to studying comparative development using iPSC models, which we have discussed, this model provides the opportunity to study previously unappreciated aspects of primate biology.

## Methods

### Human and chimpanzee iPSC panels

In this study, we include four chimpanzee iPSC lines (2 males, 2 females) from a previously described panel (*16*) and six human lines (3 males, 3 females) (*17*) matched for cell type of origin, reprogramming method, culture conditions and closely matched to passage number (median passage was within 1 passage across species and differentiation batches). We evaluated iPSC lines for pluripotency measures, differentiation potential, lack of integrations and normal karyotypes as described previously (*16, 17*) (Figure S15-17). We identified one human individual (H5) that tested positive for episomal vector sequence (Figure S17a). This individual was not an obvious outlier in any of our data (Figure 1b, 2), thus we choose to include it in our study. Original chimpanzee fibroblast samples for generation of iPSC lines were obtained from the Yerkes Primate Center under protocol 006–12. Human fibroblasts samples for generation of iPSC lines were collected under University of Chicago IRB protocol 11–0524. Feeder free iPSC cultures were initially maintained on Growth Factor Reduced Matrigel using Essential 8 Medium (E8) as previously described. After 10 passages in E8, all cell lines were transitioned to iDEAL feeder free medium that was prepared in house as specified previously (*39*). Cell culture was conducted at 37°C, 5‥ CO_2_, and atmospheric O_2_.

### Endoderm Differentiation

To produce definitive endoderm and intermediate cell types, we followed a recently published three-day protocol that systematically identified and targeted pathways involved in cell fate decisions, at critical junctures in endoderm development (*11*) with minimal modification. At 12 hours prior to initiating differentiation, iPSC lines at 70-90‥ confluence were seeded at a density of 50,000 cells/cm^2^. Basal medial for differentiations consisted of 50/50 IMDM/F12 basal media supplemented with 0.5 mg/mL human albumin, 0.7 μg/mL Insulin, 15 μg/mL holo-Transferrin and 1‥ v/v chemically defined lipid concentrate. For differentiation, basal media was supplemented with the following: day 0 to day 1 (Primitive streak induction) media included 100 ng/mL Activin A, 50 nM PI-103 (PI3K inhibitor), 2 nM CHIR99021 (Wnt agonist), days 1->2 (total of 2 media changes) media included 100 ng/mL Activin A and 250 nM LDN-193189 (BMP inhibitor). Two independent differentiation batches were performed, resulting in replicates for a subset of individuals. Each chimpanzee was replicated, while only two humans individuals were replicated across the two batches. Replicates were sex-balanced both within and across species. Cell culture was conducted at 37°C, 5‥ CO_2_, and atmospheric O_2_.

### Purity assessment using flow cytometry

Cells were dissociated using an EDTA based cell release solution, centrifuged at 200 x g for 5 minutes at 4°C and washed with PBS. Subsequently, 0.5-1 million cells were fixed and permeabilized using the Foxp3 / Transcription Factor Staining Buffer Set from eBioscience. Cells were fixed at 4°C for 30 minutes before washing once using FACS buffer (autoMACS® Running Buffer, Miltenyi Biotech). 150,000 cells were transferred to BRAND lipoGrade 96 well immunostaining plates and centrifuged at 200 x g for 5 minutes at 4°C. Cells were rinsed in FACS buffer then resuspended in the staining solution. A single master mix containing 1X Permeabilization buffer (eBioscience), BD Horizon Brilliant Stain Buffer and antibodies was prepared and 30 μL of this mix was added to each well containing cells. In order to estimate purity for each day of the timecourse, we utilized a mixture of six different directly labeled antibodies: Oct3/4 (BV421 labeled clone 3A2A20, Biolegend), SOX2 (PerCP-Cy5.5 labeled clone O30-678, BDbio), SOX17 (Alexa 488 labeled clone P7-969, BDbio), EOMES (PE-Cy7 labeled clone WD1928, eBioscience),CKIT (APC labeled clone 104D2, Biolegend), CXCR4 (BV605 labeled clone 12G5, Biolegend). All antibodies were used at the manufacturer recommended dilution except CKIT and CXCR4, which was used at 1/10 of the manufacturer specified concentration (15 ng of each antibody in final volume of 30 μL per staining). We found that the manufacturer recommended dilution produced acceptable results for live cells, however, upon fixing, we observed nonspecific binding by all populations. Thus we determined the optimal antibody titer to maximize the separation between iPSCs (biological negative) and Day 3 definitive endoderm. We found this optimal concentration to be in concordance with that quantity specified by a previous publication using the same antibody clone from a different manufacturer (39). Cells were stained for 1 hour at 4°C and subsequently washed 3x using a solution of BD Horizon Brilliant Stain Buffer containing 1X Permeabilization buffer, on the final wash cells were resuspended in 100 μL FACS buffer for acquisition on a BD LSR II flow cytometer. After data acquisition compensation, we used the program FlowJo (http://docs.flowjo.com/d2/credits-2/) to determine scaling. To do so, we used data from single stained compensation beads (Life Technologies) that were stained and collected in parallel. Live, intact, single cells were gated based on FSC and SSC channels as previously described (*11*). Day 0 iPSC purity was estimated by dual positive OCT3/4 and SOX2 (*40*) as well as negative staining for EOMES. Day 1 primitive streak purity was estimated primarily based on EOMES Positive staining (*23, 41*) but also negative staining for SOX17. Day 2 endoderm progenitor purity was quantified by positive staining for SOX17 expression (*42*) (CKIT could also be used, as its level peaks at day 2) and negative staining for CXCR4. Finally, day 3 definitive endoderm purity was estimated by double staining for CKIT and CXCR4 (*43*). For all time points, cells were stained with the full complement of markers; initial gates were defined using fluorescence intensity levels of an iPSC line as a biological negative control for days 1, 2, and 3. For day 0 (iPSCs), a definitive endoderm time point was used to quantify the biological negative for OCT3/4 and SOX2 fluorescence intensity. All iPSC lines regardless of species were at comparable fluorescence intensity levels, so we choose a representative chimp and human line to use as our standard for defining and refining all gates. Fully resolving all time points simultaneously required us to define high and low staining gates, which were determined using the time points for that marker’s maximum and minimum fluorescence intensities. All gates were refined using the same two representative chimpanzee and human lines as used for determining biological negatives, resulting in one universal gating scheme that was applied to both species and all time points. A complete gating scheme is outlined in Figure S1a-b, with the final purity results for the second batch of differentiation in Supplementary Data S1. The samples in the first differentiation batch demonstrated hallmarks of improper fixing (highly nonspecific staining of antibodies, most notably for surface markers CXCR4 and CKIT), thus we were unable to determine reliable purity estimates for the first differentiation batch.

### RNA extraction, library preparation, and sequencing

We collected RNA from iPSCs (day 0) prior to adding day 1 media, and then every 24 hours during the differentiation timecourse for a total of 4 time points representing intermediate cell populations from iPSCs to definitive endoderm (Figure S6b). We extracted the RNA using the ZR-Duet DNA/RNA MiniPrep kit (Zymo) with the addition of an on column DNAse I treatment step prior to RNA elution. To estimate the RNA concentration and quality, we used the Agilent 2100 Bioanalyzer (Figure S2). We added barcoded adaptors (Illumina TruSeq RNA Sample Preparation Kit v2) and sequenced the 50 base pair single-end RNA-seq libraries on the Illumina HiSeq 4000 at the Functional Genomics Core at University of Chicago on two flowcells (Supplementary Data S3). To minimize the introduction of biases due to batch processing, we chose the RNA extraction batches, library preparation batches, sequencing pools, adaptor names, and flowcells in a manner that maximally partitioned the biological variables of interest (day, species, cell line; Supplementary Data S3, S5).

We generated a minimum of 14,424,520 raw reads per sample. We used FastQC (http://www.bioinformatics.babraham.ac.uk/projects/fastqc/) to confirm that the reads were high quality.

### Quantifying the number of RNA-seq reads from orthologous genes

We mapped human reads to the hg19 genome and chimpanzee reads to panTro3 using TopHat2 (version 2.0.11) (*18*), allowing for up to two mismatches in each read. We kept on only reads that mapped uniquely. To prevent biases in expression level estimates due to differences in mRNA transcript size and the relatively poor annotation of the chimpanzee genome, we only kept reads that mapped to a list of orthologous metaexons across 30,030 Ensembl genes as described previously (*10*). Gene expression levels were quantified using the *feature counts* function in SubRead 1.4.4 (*44*). For one sample (C2B at Day 0), the number of raw reads was approximately double the second highest number of raw reads. Therefore, we subsampled the raw reads to approximately the same number of raw reads as the second highest sample.

We performed all downstream processing and analysis steps in R (version 3.2.2) unless otherwise stated.

### Transformation and normalization of RNA-sequencing reads

After receiving the raw gene counts, we calculated the log_2_-transformed counts per million (CPM) for each sample using edgeR (*21*). To filter for the lowly expressed genes, we kept only genes with an expression level of log_2_(CPM) > 1.5 in at least 16 samples per species (*45*). For the remaining genes, we normalized the original read counts using the weighted trimmed mean of M-values algorithm (TMM) (*45*) to account for differences in the read counts at the extremes of the distribution and calculated the TMM-normalized log_2_-transformed CPM.

When we performed principal components analysis (PCA) using the TMM-normalized log_2_-transformed log2(CPM) values, we found one outlier (H1B at Day 0, Figures S3a). We removed this sample from the list of original gene counts. We filtered for the lowly expressed genes by retaining genes with an expression level of log_2_(CPM) > 1.5 in at least 15 human samples and at least 16 chimpanzee samples. 10,304 genes remained. We performed TMM-normalization and then performed a cyclic loess normalization with the function normalizeCyclicLoess from the R/Bioconductor package limma (*20, 46*). We found that the TMM-normalized log_2_(CPM) values were highly correlated with the TMM-and cyclic loess-normalized log_2_(CPM) values (r > 0.99 in the 63 samples). We used the TMM-and cyclic loess-normalized log_2_(CPM) expression values in all downstream analysis unless otherwise stated.

We calculated normalized log_2_-transformed RPKM values by using the function rpkm with normalized library sizes from the package edgeR (*21*) (Figure S4b). We measured the “gene lengths” as the sum of the lengths of the orthologous exons and were also used in (*16*). Our RPKM calculation is robust to the calculation method. This method of calculating RPKM was highly correlated with a method in which we subtracted log_2_(gene length in kbp) from the TMM-and cyclic loess-normalized log_2_(CPM) values (r > 0.97).

### Data quality and analysis of technical factors

To assess the data quality, we performed Principal Components Analysis (PCA) on the normalized log_2_(CPM) values from above (Figure 2a). Principal component (PC) 1 was highly associated with day and PC2 was highly associated with species (r > 0.92 for each, Figure 2a; Supplementary Data S5). We sought to determine if the study’s biological variables of interest were confounded with any of the study’s recorded technical aspects (Supplementary Data S1-3). First, we calculated which of our 35 recorded technical factors were statistically significant predictors of PCs 1-5 with individual linear models for each technical factor. The 19 statistically significant predictors (FDR cutoff of 10‥ assessed on the 5x35 matrix) were carried to the second stage. In this stage, we determined which technical factors were associated specifically with either day or species, with individual linear models for each technical factor. We quantified these associations using the *P* values from analysis of variance (ANOVA) for the numerical technical factors and from Chi-squared test (using Monte Carlo simulated *P* values) for the categorical technical factors. Statistical significance was determined by Benjamini-Hochberg adjusted *P* value

< 10‥ (assessed on the 2x19 matrix). Variables for cell line and sex-by-species include a species but not a day component and were tested in this pipeline. They were found to each be confounded with species (B.H. adj. *P* value = 0.0095 each) but not day, thereby increasing the confidence in our pipeline.

During this analysis, we observed that the purity estimates were relatively similar across days and between species until the final day (Figure S1d; Supplementary Data S1). Therefore, it was important to explore how the variance for a given technical factor was partitioned across the biological variables of interest (e.g. across the days, species, and day-by-species interactions). For each recorded technical variable, we created a reduced model and a full model. The reduced model contained only species and day as fixed effects and the technical factor as the response variable. The full model had the same response variable but contained species, day, and a species-by-day interaction as fixed effects. We then compared the two models and reported the significance (Supplementary Data S13).

The exact tools used to compare the two models were data-dependent (Supplementary Data S3 and columns 1-2 in Supplementary Data S13). For numerical data (24 technical factors), we constructed the full and reduced normal general linear models for each technical factor. We compared the models using ANOVA, and extracted the *P* value directly from ANOVA. For categorical data with 2 levels (3 technical factors), we constructed the two general linear models from the binomial family. We used ANOVA to compare the models and extracted the deviance along with its degrees of freedom. Based on the deviance, we calculated the Chi-Squared statistic and associated *P* value. 8 technical factors (such as RNA extraction data) contained categorical data with more than two levels. We modeled this data type with multinomial logistic regression with the R/Bioconductor package nnet (*47*) and used ANOVA to obtain the likelihood ratio statistic and associated *P* value. We performed this process for each technical factor using data from days 0 and 1 as well as from days 0 to 3 (Supplementary Data S13).

### A linear model based framework to perform pairwise differential expression analysis

Differential expression was estimated using a linear model based empirical Bayes method implemented in the R package limma (*48, 49*). In order to use a linear modeling approach with RNA-seq read counts, we calculated weights that account for the mean-variance relationship of the count data using the function voom from the limma package (*50*). This limma+voom pipeline has previously been shown to perform well with n > 3 biological replicates/condition (*51, 52*).

For all pairwise differential expression comparisons, the species, day, and a species-by-day interaction were modeled as fixed effects, and individual as a random effect. Individual (cell line) rather than differentiation batch was modeled as a random effect because when using a linear model, individual was most highly correlated with PCs 2 and 3, whereas batch was most highly correlated with PC 10. Since our recorded technical factors were not confounded with our biological variables of interest and did not contribute significantly to the first five principle components of variation (Supplementary Data S5), we did not include any other covariates. (See Discussion regarding variables inherent to iPSCs and iPSC-differentiated cells.) We used contrast tests in limma to find genes that were differentially expressed (DE) by species at each day (Supplementary Data S6), DE between days for each species (Supplementary Data S7), and significant day-by-species interactions for days 1-3 (Supplementary Data S9). For each pairwise DE test, we corrected for multiple testing with the Benjamini & Hochberg false discovery rate (*53*) and genes with an FDR-adjusted *P* values <0.05 were considered DE unless otherwise stated in the text.

To find the number of shared DE genes in consecutive time points in each species (Figure 4), we used a two *P* value cutoff system. To be “shared” across species for a given pair of time points (e.g. day 0 to 1), a gene must have an FDR-adjusted *P* value <0.01 in one species and an FDR-adjusted *P*-value <0.05 in the other species (*53*). To estimate the percentage of DE genes in chimpanzees given the observation in humans, we divided the number of genes with an FDR-adjusted *P* value <0.01 in chimpanzees over the number of genes with an FDR-adjusted *P* value <0.05 in humans.

### Combining technical replicates

Some analyses did not allow us to model technical replicates explicitly (and treating them as biological replicates would introduce bias in the data). Therefore, we combined technical replicates for the same individual, when available. We calculated the average of the normalized log_2_(CPM) values for each cell line at each time point. For day 0, 1 human cell line had a pair of technical replicates that were averaged together. For days 1-3, 2 human cell lines had technical replicates that were averaged. We were able to average technical replicates for each of the four chimpanzee cell lines at each time point. After this process, 6 human data points and 4 chimpanzee data points per day remained, for a total of 40 data points.

When we performed principal components analysis (PCA) using these 40 data points, the results were similar to the PCA plot including all the technical replicates (Figure S7a)-PC1 was still correlated with day and PC2 was correlated with species (Supplementary Data S5). We visually inspected the PCA plot for the distinct clustering of data points with averaged technical replicates and single replicates in the humans, and this potential pattern was not present, increasing our confidence that this process did not introduce bias into the data.

We found that the expression values for the 40 samples were robust with respect to the method used to combine the technical replicates. The post-normalization method described above was strongly correlated with a pre-normalization method to combine technical replicates (r > 0.99 for the 10,304 genes included in the main analysis; Figure S7b). In our pre-normalization method of combining the technical replicates, we summed the raw counts of technical replicates at each time point (for a total 40 data points) and performed the normalization steps described in the “Counting and Normalization” section.

### Joint Bayesian analysis with Cormotif

To cluster genes by their temporal gene expression patterns, we used the R/Bioconductor package Cormotif (version 1.18.0), a method that jointly models multiple pairwise differential expression tests (*25*). Unlike other available methods in this class, the Cormotif framework allows for data-set specific differential expression patterns. To identify patterns in expression over time (called “correlation motifs”), expression levels from days 1-3 were compared to those to the previous day for each gene in each species. Since the program does not allow for the explicit modeling of technical replicates (unlike the voom+limma method above), we first ran the program with the expression values averaged across technical replicates. For more information on this process, see the Methods section on “Combining technical replicates”.

To use Cormotif, we were required to specify the number of correlation motifs to model. We determined a reasonable range by investigating both the Bayesian information criterion (BIC) and Akaike information criterion (AIC). We observed that the BIC and AIC were minimized across many seeds when 7 or 8 correlation motifs were modeled, respectively (Figure S8a). Thus we further explored models with 7 and 8 correlation motifs. Because Cormotif is not deterministic, we ran Cormotif 100 times and recorded the seed that produced the model with the largest log likelihood (Supplementary Data S16). The best model (the seed with the greatest log likelihood) with 7 correlation motifs is displayed in Figure S8b, and the best model with 8 correlation motifs is featured in Figure 5. We selected the model with 8 correlation motifs to be the primary figure because it had a large log likelihood and all motifs contained more than 100 genes. It should be noted, however, that the two models had very similar correlation motifs (expression patterns; Supplementary Information).

We were initially conservative when assigning a gene to a specific correlation motif. Following the advice of the Cormotif authors (*25*), a gene must have a posterior likelihood estimate of ≥ 0.5 to be called DE between time points and < 0.5 to be considered not DE. We also used this assignment criteria when using Cormotif to compare expression levels using different combination methods (Figures S8c) and to compare all time points to day 0 (Figure S8d). For a trajectory to be defined as DE, the trajectory in humans and chimpanzees needed similar posterior probabilities of differential expression (≤ 0.20) at each comparison along the trajectory. We performed GO enrichment analysis (*27, 28*) on various combinations of correlation motifs (Supplementary Data S10, 11) using the PANTHER Overrepresentation Test tool (release 20160715) from the Panther Database (*26, 54*) (http://pantherdb.org/tools/compareToRefList.jsp).

### Global analysis of variation in gene expression levels

We calculated the variance in gene expression level for each gene in each species. Since the largest theoretical range of a variance is from 0 to infinity, we performed a log_2_ transformation to each variance value. We then performed a one-sided t-test between the distribution of log_2_(variances in gene expression) from day 0 and day 1 in each species, with the alternative hypothesis that the variation was greater in day 0 than day 1. We compared the effect sizes of interspecies DE genes with a one-sided Mann–Whitney *U* test on magnitudes of effect sizes (Figure S10). We tested the null that there was no change in log_2_ fold change in gene expression across the species from day 0 to day 1, with the alternative hypothesis that the average magnitude of effect of DE genes (FDR = 5‥) was greater in day 0 than day 1.

### Gene-by-gene analysis of variation in gene expression levels and calculating the proportion of true positives

To determine if there was an enrichment of genes undergoing changes in variation one species, we used an F test to compare two variances in R (var.test command) for each gene using the averaged log_2_(CPM) expression values of technical replicates. In these tests, the null hypothesis was no change of variance in the gene expression levels between days and the alternative hypothesis was a reduction in variation of gene expression levels between two time points (a one-sided test).

We calculated the *P* values for the F statistics from each test and plotted the densities using ggplot2 (*55*). If a *P* value distribution appeared to be even slightly skewed towards small *P* values, we used the R package qvalue (https://github.com/StoreyLab/qvalue) to determine Πo, the true proportion of null statistics from a given *P* value distribution (*30*). Its complement, Π1, is considered the proportion of significant tests from a *P* value distribution. We used this process to analyze the reduction in variation in each species from days 0 to 1, 1 to 2, 2 to 3, and 0 to 2 (Figure 7a-b).

Afterwards, we used the same procedure (F tests) to test the alternative hypothesis that the variation of gene expression increased between two time points. We determined Πo and Π1 in the same manner as above to analyze the increase in variation in each species from days 0 to 1, 1 to 2, 2 to 3, and 0 to 2 (Figure 7a-b).

### Estimating the proportion of genes that undergo a change in variation in both species

We then estimated the true proportion of significant genes shared across species for a given set of time points. Rather than take the intersection of the significant genes (for which we would be underpowered), we adopted a method from Storey and Tibshirani 2003 (Storey’s Πo) (*30*). This method was recently implemented by Banovich 2016 to determine the sharing of quantitative trait loci (QTLs) from different cell types (*29, 30*). Using the *P* value distributions generated in the previous section, we subset the genes in species 2 conditioned on its F statistic significance in species 1 (unadjusted *P* value < 0.05). To test for an enrichment of small *P* values, we used the *P* values from species 2 to determine Πousing the same process as the previous section (Figure 7c-f, S13). We then repeated this process for other *P* value cutoffs, including 0.01 and 0.10 (Figure S14e-l). To determine robustness with respect to the number of genes considered significant in species 1, we calculated Πo for species 2 conditioned on 100 genes with the lowest *P* values in species 1. This process was repeated for the top 101 to all 10,304 genes (Figure S14a-d).

### Estimating the null hypothesis for the proportion of genes that undergo a change in variation in both species

To determine the null hypothesis for the Π1 based on conditioning, we performed permutation tests. First, we combined the unadjusted *P* values from the F test for a reduction in variation from days 0 to 1 in chimpanzees (species 1) and humans (species 2). We used the randomizeMatrix function in the R package picante (*56*) to permute the *P* values of species 1 and then merged this *P* value distribution with the *P* value distribution from species 2. We then determined Πo in species 1 conditioned on its *P* value significance in species 2 (unadjusted *P* value < 0.05). We repeated this process a total of 100,000 times and found the complement of the 100,000 values (Supplementary Data S15). We defined the permuted null hypothesis as the mean Π1 value. We then repeated this process, with humans as species 1 and chimpanzees as species 2.

### Data Access

The data have been deposited in NCBI’s Gene Expression Omnibus (*57*) and are accessible through GEO Series accession number GSE98411. The website for this dataset is http://www.ncbi.nlm.nih.gov/geo/query/acc.cgi?acc=GSE98411. Our custom pre-processing script is available upon request to the authors.

All data and scripts used post-processing are available at https://github.com/Lauren-Blake/Endoderm_TC/tree/gh-pages. All of our analyses using R and the subsequent results can be viewed at https://lauren-blake.github.io/Endoderm_TC/analysis/.

## Acknowledgment

We thank members of the Gilad, Stephens, and Pritchard labs for helpful discussions and comments on the manuscript. L.E.B. was supported by the National Science Foundation Graduate Research Fellowship (DGE-1144082) and by the Genetics and Gene Regulation Training Grant (T32 GM07197). B.J.P. was supported by the Training in Emerging Multidisciplinary Approaches to Mental Health and Disease (T32MH020065). This work was funded by an NIH grant from NIGMS (MH084703). The content presented in this article is solely the responsibility of the authors and does not necessarily reflect the official views of the National Institutes of Health or the National Science Foundation.

## Author contributions

Y.G. and B.J.P. conceived of the study and designed the experiments. S.T. and B.J.P. performed the experiments. M.M. and C.C. extracted the RNA and prepared the sequencing libraries. L.E.B., S.T., and B.J.P. analyzed the results (with input from J.D.B. and C.J.H.). Y.G. and B.J.P. supervised the project. L.E.B., S.T., Y.G., and B.J.P wrote the paper.

